# How hot is the hot zone? Computational modelling clarifies the role of parietal and frontoparietal connectivity during anaesthetic-induced loss of consciousness

**DOI:** 10.1101/2020.12.19.423595

**Authors:** Riku Ihalainen, Olivia Gosseries, Frederik Van de Steen, Federico Raimondo, Rajanikant Panda, Vincent Bonhomme, Daniele Marinazzo, Howard Bowman, Steven Laureys, Srivas Chennu

## Abstract

In recent years, specific cortical networks have been proposed to be crucial for sustaining consciousness, including the posterior hot zone and frontoparietal resting state networks (RSN). Here, we computationally evaluate the relative contributions of three RSNs – the default mode network (DMN), the salience network (SAL), and the central executive network (CEN) – to consciousness and its loss during propofol anaesthesia. Specifically, we use dynamic causal modelling (DCM) of 10 minutes of high-density EEG recordings (*N* = 10, 4 males) obtained during behavioural responsiveness, unconsciousness and post-anaesthetic recovery to characterise differences in effective connectivity within frontal areas, the posterior “hot zone”, frontoparietal connections, and between-RSN connections. We estimate – for the first time – a large DCM model (LAR) of resting EEG, combining the three RSNs into a rich club of interconnectivity. Consistent with the hot zone theory, our findings demonstrate reductions in inter-RSN connectivity in the parietal cortex. Within the DMN itself, the strongest reductions are in feed-forward frontoparietal and parietal connections at the precuneus node. Within the SAL and CEN, loss of consciousness generates small increases in bidirectional connectivity. Using novel DCM leave-one-out cross-validation, we show that the most consistent out-of-sample predictions of the state of consciousness come from a key set of frontoparietal connections. This finding also generalises to unseen data collected during post-anaesthetic recovery. Our findings provide new, computational evidence for the importance of the posterior hot zone in explaining the loss of consciousness, highlighting also the distinct role of frontoparietal connectivity in underpinning conscious responsiveness, and consequently, suggest a dissociation between the mechanisms most prominently associated with explaining the contrast between conscious awareness and unconsciousness, and those maintaining consciousness.

**Highlights:**

- Modelling shows that connectivity within hot zone tracks change of conscious state
- Separately, frontoparietal connections support maintenance of conscious state
- Strength of frontoparietal connections predicts conscious state in unseen data
- Both parietal hot zone and frontoparietal connectivity important for consciousness

**Funding:** This work was supported by the UK Engineering and Physical Sciences Research Council (EP/P033199/1), Belgian National Funds for Scientific Research (FRS-FNRS), the University and University Hospital of Liege, the Fund Generet, the King Baudouin Foundation, the AstraZeneca Foundation, the European Union’s Horizon 2020 Framework Programme for Research and Innovation under the Specific Grant Agreement No. 945539 (Human Brain Project SGA3), DOCMA project (EU-H2020-MSCA–RISE–778234), the BIAL Foundation, the European Space Agency (ESA) and the Belgian Federal Science Policy Office (BELSPO) in the framework of the PRODEX Programme, the Center-TBI project (FP7-HEALTH-602150), the Public Utility Foundation ‘Université Européenne du Travail’, “Fondazione Europea di Ricerca Biomedica”, the Mind Science Foundation, the European Commission, and the Special Research Fund of Ghent University. O.G. is research associate and S.L. is research director at the F.R.S-FNRS.

**Declaration of interest:** None.

**Significance Statement:** Various connectivity studies have suggested multiple network-level mechanisms driving changes in the state of consciousness, such as the posterior hot zone, frontal-, and large-scale frontoparietal networks. Here, we computationally evaluate evidence for these mechanisms using dynamic causal modeling for resting EEG recorded before and during propofol-anaesthesia, and demonstrate that, particularly, connectivity in the posterior hot zone is impaired during propofol-induced unconsciousness. With a robust cross-validation paradigm, we show that connectivity in the large-scale frontoparietal networks can consistently predict the state of consciousness and further generalise these findings to an unseen state of recovery. These results suggest a dissociation between the mechanisms most prominently associated with explaining the contrast between conscious awareness and unconsciousness, and those maintaining consciousness.

## 1. Introduction

Several cortical network-level mechanisms have been proposed to explain human consciousness and its loss, of which two, in particular, have received an increasing amount of interest and evidence. On the one hand, empirical studies have suggested that the loss of consciousness (LOC)^1^ is associated with disruptions of within- and between-network connectivity in cortical areas associated with large-scale frontoparietal networks (Bor & Seth, 2012; Laureys & Schiff, 2012). On the other, temporo-parieto-occipital areas – colloquially named as ‘the posterior hot zone’ – has been shown to be important in mediating changes in consciousness during sleep (Siclari et al., 2017; Lee et al., 2019), and in patients with brain damage (Vanhaudenhuyse et al., 2010; Wu et al., 2015).

In this context, general anaesthetics are a powerful tool to investigate alterations in brain connectivity during changes in the state of consciousness (see Bonhomme et al., 2019 for a recent review). Indeed, several previous studies have utilised anaesthetic drugs in investigating brain dynamics in both functional and effective/directed connectivity studies and suggested multiple explanatory mechanisms of the LOC. Note that here, effective connectivity is defined following (Friston, 2011) and (Razi & Friston, 2016) as a causal influence (in a control theory sense) of one neural population over another and functional connectivity as undirected statistical dependencies between distinct neurophysiological events. Some of these studies have suggested a breakdown of thalamo-cortical connections and disrupted frontoparietal networks (Boveroux et al., 2010; Schrouff et al., 2011). Others have found disruptions in frontal areas (Guldenmund et al., 2016), diminished frontoparietal feedback connectivity (Lee et al., 2009; Lee, Ku et al., 2015), and increased frontoparietal connectivity (Barrett et al., 2012). To bring computational evidence to bear upon this discussion, we adopt one of the most commonly used methods for understanding effective connectivity, dynamic causal modeling (DCM; Friston, Harrison & Penny, 2003), to assess cortical network-level mechanisms involved in the LOC, and evaluate the evidence for the posterior hot zone.

There are relatively few studies assessing resting state effective connectivity with DCM during anaesthetic-induced unconsciousness, but a recent fMRI study identified impaired subcortico-cortical connectivity between globus pallidus and posterior cingulate (PCC) nodes, but no cortico-cortical modulations (Crone, Lutkenhoff, Bio, Laureys, & Monti, 2017). Boly et al. (2012) found a decrease in feedback connectivity from frontal (dorsal anterior cingulate; dACC) to parietal (PCC) nodes. Both of these studies, however, evaluated relatively simple models in terms of cortical sources (excluding subcortical nodes), consisting of only two such nodes – an anterior and a posterior node. Consequently, they do not allow us to compare the role of the posterior hot zone to other potential cortical mechanisms underpinning consciousness.

Here, we address this gap by modelling changes in key resting state networks (RSN) - the default mode network (DMN), the salience network (SAL), and the central executive network (CEN), due to unconsciousness induced by propofol, a common clinical anaesthetic. We employ a novel methodological combination of DCM for resting EEG cross-spectral densities (CSD; Friston et al., 2012; Moran et al., 2009) and Parametric Empirical Bayes (PEB; Friston et al., 2016), to better estimate model parameters (and their distributions) and prune redundant connections. Within this framework, we invert - for the first time - a single large-scale model of EEG, consisting of 14 RSN nodes, in addition to the individual RSNs themselves (figure 1). This allows us to evaluate the role of different subgroups of intra- and inter-RSN connections in the modulation of consciousness. Further, we apply robust leave-one-subject-out-cross-validation (LOSOCV) on DCM model parameters, to evaluate hypotheses about whether specific sets of connections within and between frontal and parietal nodes are not only able to explain changes between states of consciousness, but also to predict the state of consciousness from unseen EEG data. Using this combination of computational modelling, cross-validation and hypothesis testing, we indicate the importance of the posterior hot zone in explaining the loss of consciousness, while highlighting also the distinct role of frontoparietal connectivity in underpinning conscious responsiveness. Consequently, we demonstrate a dissociation between the mechanisms most prominently associated with explaining the contrast between conscious awareness and unconsciousness, and those maintaining consciousness.

**Figure 1.**
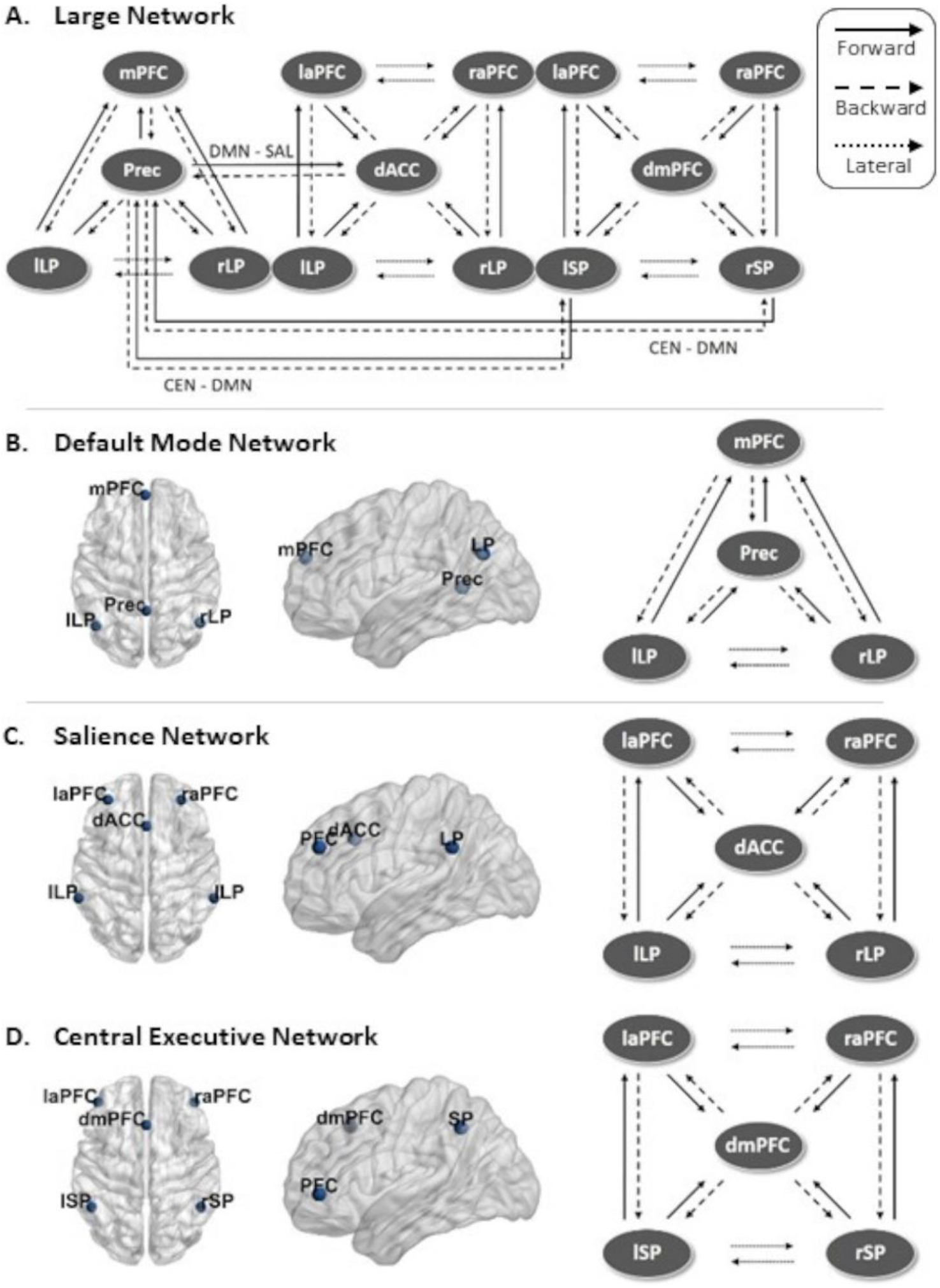
Full model schematics and node locations. **A.** Schematic view of the large DCM model consisting of the 14 nodes and connections combining three RSNs. Inter-RSN connections were specified between PCC/precuneus and bi-lateral superior parietal nodes, and between PCC/precuneus and anterior cingulate cortex. **B-D.** Location of the nodes and the schematic representation of the full model for DMN, SAL, and CEN, respectively.

## 2. Methods

### 2.1 Data acquisition and preprocessing

The data used in the present work were acquired from a previous propofol anaesthesia study, which describes the experimental design and data collection procedure in detail (Murphy et al., 2011). The study was approved by the Ethics Committee of the Faculty of Medicine of the University of Liège, and written consent was obtained from all the participants. None of the participants suffered from mental illness, drug addiction, asthma, motion sickness, nor had a history of mental illness or suffered from any previous problems with anaesthesia. The data consisted of 15 minutes of spontaneous, eyes-closed high-density EEG recordings (256 channels, EGI) from 10 participants (mean age 22 ± 2 years, 4 males) in four different states of consciousness: behavioural responsiveness, sedation (Ramsay scale score 3, slower responses to command), loss of consciousness with clinical unconsciousness (Ramsay scale score 5-6, no response to command), and recovery of consciousness (Ramsay, Savege, Simpson, & Goodwin, 1974). Note that for the recovery state, the data consisted of 9 datasets. Participants were considered to be fully awake if the response to verbal command (‘squeeze my hand’) was clear and strong (Ramsay 2), and in LOC, if there was no response (Ramsay 5-6). The Ramsay scale verbal commands were repeated twice at each level of consciousness. Propofol was infused through an intravenous catheter placed into a vein of the right hand or forearm, and the propofol plasma and effect-site concentrations were estimated with 3.87 ± 1.39 mcg/mL average arterial blood concentration of propofol for LOC. Here, we only modelled data from the maximally different anaesthetic states, behavioural responsiveness and LOC, and used recovery as a test of DCM model generalisation. These data can be made available after signing a formal data-sharing agreement with the University of Liège.

Data from channels from the neck, cheeks, and forehead were discarded as they contributed most of the movement-related noise, leaving 173 channels on the scalp for the analysis. These 173 electrodes were co-registered to a template MRI mesh in MNI coordinates, and the volume conduction model of the head was based on the Boundary Element Method (BEM). The raw EEG signals were filtered from 0.5 – 45 Hz with additional line noise removal at 50 Hz using a notch filter. The recordings were then downsampled to 250 Hz, and abnormally noisy channels and epochs were identified by calculating their normalised variance, and then manually rejected or retained by visual inspection. Last, the data were then re-referenced using the average reference.

### 2.2 Dynamic causal modeling

For the DCM modelling of the high-density EEG data, the first 60 artefact-free 10-second epochs in wakeful behavioural responsiveness and LOC were combined into one dataset with two anaesthetic states making up a total of 120 epochs per participant. The preprocessed data was imported in to SPM12 (Wellcome Trust Centre for Human Neuroimaging; www.fil.ion.ucl.ac.uk/spm/software/spm12).

To analyse effective connectivity within the brain’s resting state networks, DCM for EEG cross-spectral densities (CSD) was applied (Friston et al., 2012; Moran et al., 2009). Briefly, with this method, the observed cross-spectral densities in the EEG data are explained by a generative model that combines a biologically plausible neural mass model with an electrophysiological forward model mapping the underlying neural states to the observed data. Each node in the proposed DCM models – that is, each electromagnetic source – consists of three neural subpopulations, each loosely associated with a specific cortical layer; pyramidal cells, inhibitory interneurons and spiny stellate cells (ERP model; Moran, Pinotsis & Friston, 2013). DCM does not simply estimate the activity at a particular source at a particular point in time – instead, the idea is to model the source activity over time, in terms of interacting inhibitory and excitatory populations of neurons.^2^

The subpopulations within each node are connected to each other via *intrinsic* connections, while nodes are connected to each other via *extrinsic* connections. Three types of extrinsic connections are defined, each differing in terms of their origin and target layers/subpopulation: forward connections targeting spiny stellate cells in the granular layer, backward connections targeting pyramidal cells and inhibitory interneurons in both supra- and infragranular layers, and lateral connections targeting all subpopulations. This laminar specificity in the extrinsic cortical connections partly defines the hierarchical organisation in the brain. Generally speaking, the backward connections are thought to have more inhibitory and largely modulatory effect in the nodes they target (top-down connections), while forward connections are viewed as having a strong driving effect (bottom-up; Salin & Bullier, 1995; Sherman & Guillery, 1998).

The dynamics of hidden states in each node are described by second-order differential equations which depend on both, the parametrised intrinsic and extrinsic connection strengths. This enables the computation of the linear mapping from the endogenous neuronal fluctuations to the EEG sensor spectral densities, and consequently, enables the modelling of differences in the spectra due to changes in the underlying parameters; for example, the intrinsic and extrinsic connections. Here, for straight-forward interpretability, we modelled changes in extrinsic connections as a result of changes in the state of consciousness.

### 2.3 Model specification

Fitting a DCM model requires the specification of the anatomical locations of the nodes/sources a priori. Here, we modelled three canonical RSNs associated with consciousness (see for example Boly et al., 2008; Heine et al., 2012), namely the Default Mode Network (DMN), the Salience Network (SAL), and the Central Executive Network (CEN). In addition, we modelled a fourth large-scale network (LAR) combining all the nodes and connections in the three RSNs above, with additional inter-RSN connections motivated by structural connectivity (details below). The node locations of the three RSNs modelled here were taken from Razi et al. (2017) and are shown in figure 1 with their respective schematic representations (the node locations in figure 1 and the effective connectivity modulations in figures 4A, 5A, 6A, and 7A were visualized with the BrainNet Viewer (Xia, Wang, & He, 2013, http://www.nitrc.org/projects/bnv/). The MNI coordinates are listed in table 1. Coincidentally, these same data have been previously source localised to the same locations as some of the key nodes in the RSNs modelled here (Murphy et al., 2011). We treated each node as a patch on the cortical surface for constructing the forward model (‘IMG’ option in SPM12; Daunizeau, Kiebel, & Friston, 2009).

**Table 1.**
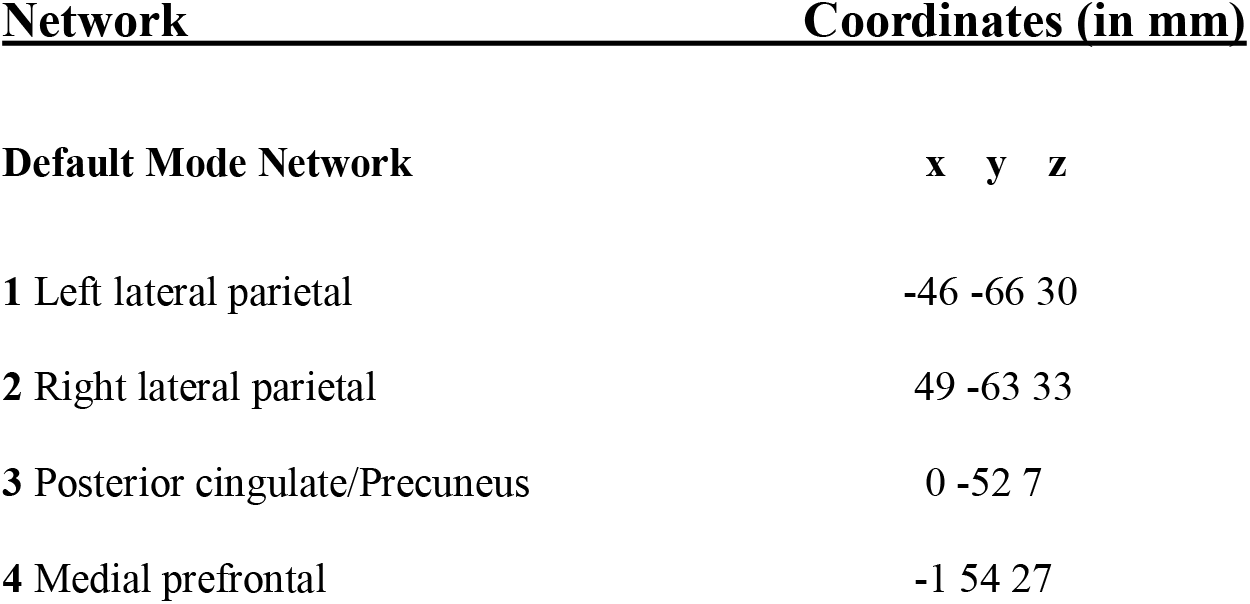
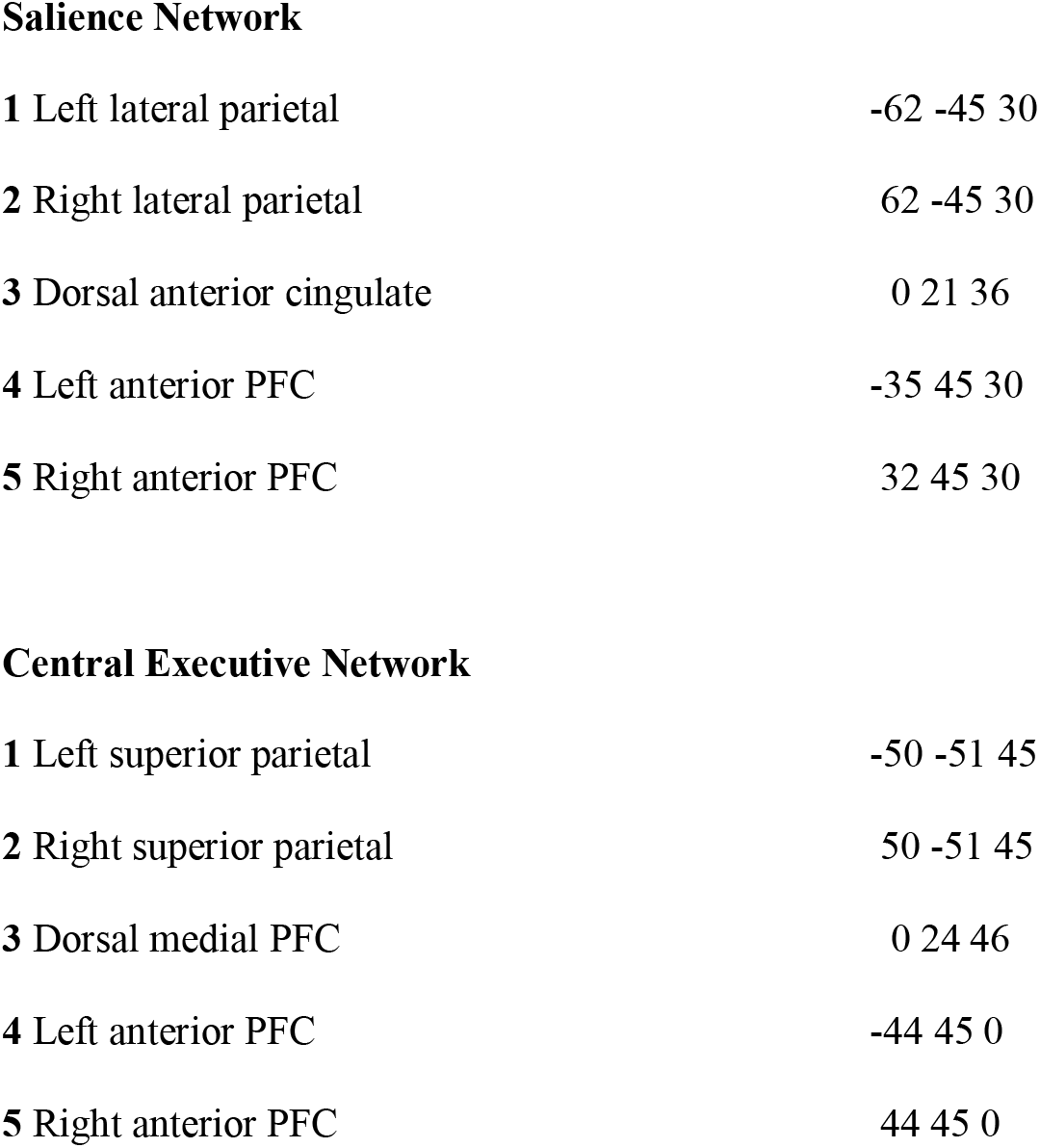
All the nodes and their corresponding MNI coordinates for the three resting state networks (adapted from Razi et al., 2017). The large model incorporated all these nodes as a single model.

Nodes in the 3 RSNs were connected via forward, backward, and lateral connections as described in David et al. (2006, 2005). Thus, each node (in each RSN-model) were modelled as a point source with the neuronal activity being controlled by operations following the Jansen-Rit model (Jansen & Rit, 1995). Note that all our models were fully connected. In addition to preserving the connections within the nodes of the original 3 RSNs, in the LAR, we additionally hypothesised potential connections between the 3 RSNs. Previous structural connectivity studies have identified a highly interconnected network of RSN hubs that seem to play a crucial role in integrating information in the brain, often termed the ‘rich-club’ (van den Heuvel & Sporns, 2011). Specifically, van den Heuvel and colleagues localised a number of these key-hubs to regions comprising of the precuneus, superior lateral parietal cortices, and superior frontal cortex, thus, to some extent overlapping with some of the key-nodes in our RSN models. Therefore, as a structurally-informed way to investigate the potential anaesthesia-induced modulations of effective connectivity between the 3 RSNs, we specified – in addition to the already-specified connections in our RSNs – bi-directional connections between PCC/precuneus and left/right superior parietal nodes (connecting DMN and CEN), and between PCC/precuneus and anterior cingulate cortex (connecting DMN and SAL).

These three different types of connections in each model were specified in what is referred in the DCM literature as the ‘A-matrix’. In addition, to explicitly parameterise the effect of the session – i.e. the effect of the anaesthetic – on the connections, we allowed every connection to change (specified in the ‘B-matrix’).

### 2.4 Model inversion

In DCM, model inversion refers to fitting the models to best explain the empirical data of each participant’s dataset, and thereby inferring a full probability density over the possible values of model parameters (with the expected values and covariance). Here, we first modelled the effects of propofol in terms of changes in connectivity that explained the differences in the empirical data observed in LOC as compared to behavioural responsiveness baseline (figure 3A). The EEG data used contained considerable peaks at the alpha range (8-12 Hz), and the default parameter settings in DCM for CSD failed to produce satisfactory fits to these peaks when inspected visually (see van Wijk et al., 2018, *p.* 824). To address this issue, we doubled the number of maximum iterations to 256 and estimated the models with two adjustments to the hyperparameters: first, we set the shape of the neural innovations (i.e. the baseline neuronal activity) to flat (−32) instead of the default mixture of white and pink (1/f) components (Moran et al., 2009). Second, we increased the noise precision value from 8 to 12 to bias the inversion process towards accuracy over complexity (see Friston et al., 2012 and Moran et al., 2009 for a detailed description of DCM for cross-spectral densities). In addition, for LAR the number of spatial modes was increased to 14 instead of the default of 8. The modes here refer to a reduction of the dimensionality of the data (done for computational efficiency) by projecting the data onto the principal components of the prior covariance, such that a maximum amount of information is retained (David et al., 2006; Fastenrath, Friston, & Kiebel, 2009; Kiebel, Garrido, Moran, & Friston, 2008).

These adjustments led to our full models (i.e. DMN, SAL, CEN, and LAR) converging with satisfactory fits (inspected visually) to the spectrum for 30/40 subject model instances (similar fits to what can be seen as the end result in figure 2). We then applied Bayesian Parameter Averaging (BPA) for each of the full models separately, averaging over the posteriors from the subject model instances that did converge and setting these averaged posteriors as new priors for the respective non-converged subject model instances. Estimating these subject model instances again with these BPA-derived priors produced satisfactory fits for all 10 remaining instances. Finally, we estimated all the full models again for all the participants with setting the posteriors from the earlier subject model estimations as updated priors, but this time with the neural innovations and noise precision set back to default settings. In doing so, all the models produced satisfactory fits with the default parameter settings for all of the participants (see figure 2).

**Figure 2.**
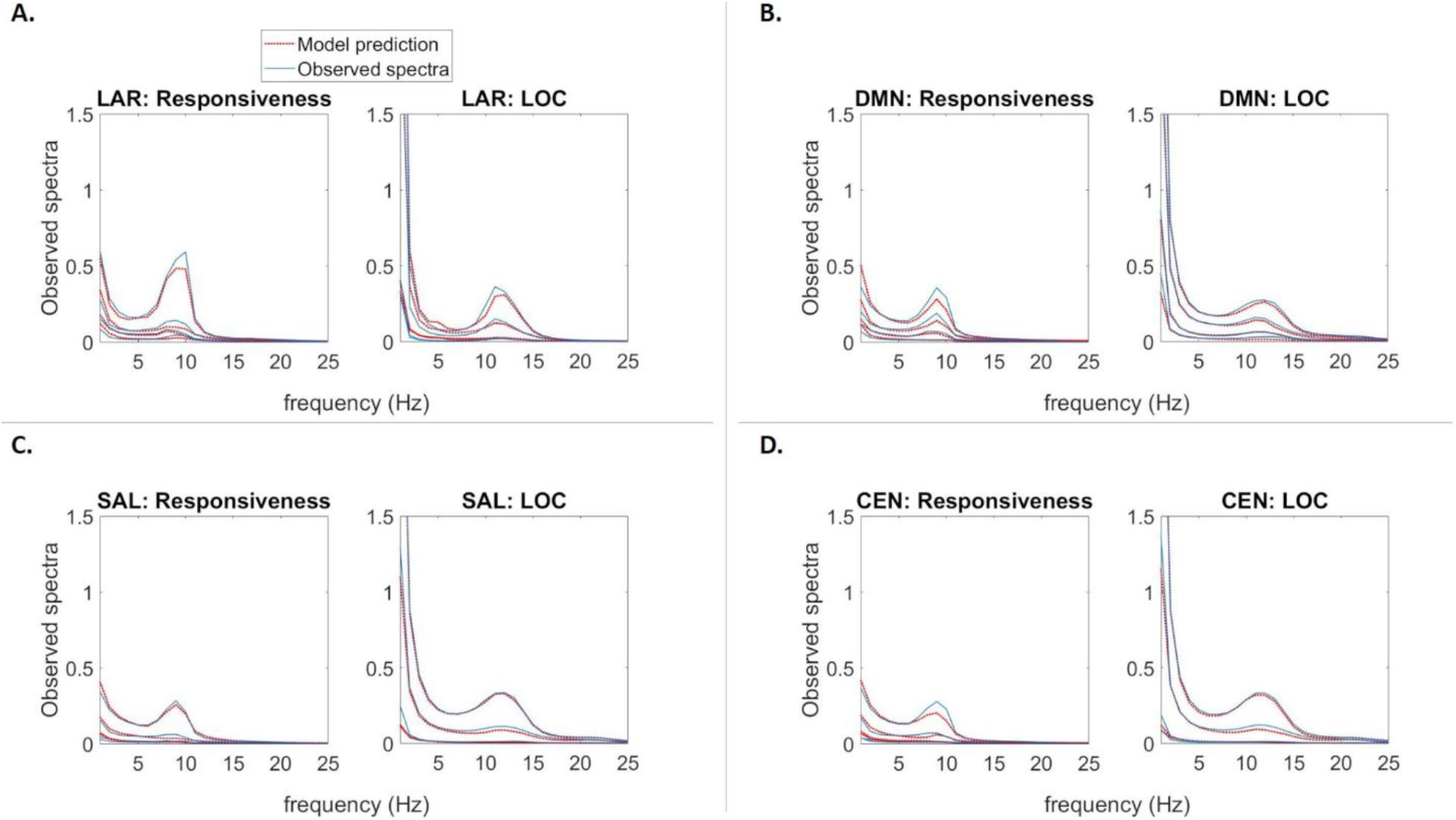
Average model fits. **A-D.** Subject-averaged power spectra of the observed EEG channel-space data, juxtaposed with that predicted by the fitted DCM models of each RSN, in normal behavioural responsiveness and LOC. Individual lines reflect spatial modes.

To validate that the priors we used in the final inversion were suitable, we compared the group-level model evidence obtained with and without the adjusted noise levels. With all full models, the default hyperparameter settings with the updated priors generated better model evidence (difference in free energies for LAR, DMN, SAL, and CEN were +47260, +9440, +15700, and +660, respectively). To qualitatively assess the model fits, the observed and model-predicted cross-spectra were visually compared in each participant and judged to be sufficiently similar. To be sure about our conclusions, we also performed the PEB modelling (see below) leaving out the fitted subject model instances that produced the worst fits (1-2 per model); this had no notable influence on the interpretation of the results. The same approach was followed when inverting the full models separately for individual states of consciousness (figure 3B); in addition to the full models, here the BPA was also restricted to the same state of consciousness. The model-predicted and original spectral densities averaged over participants are shown in figure 2A, B, C, and D for LAR, DMN, SAL, and CEN, respectively.

**Figure 3.**
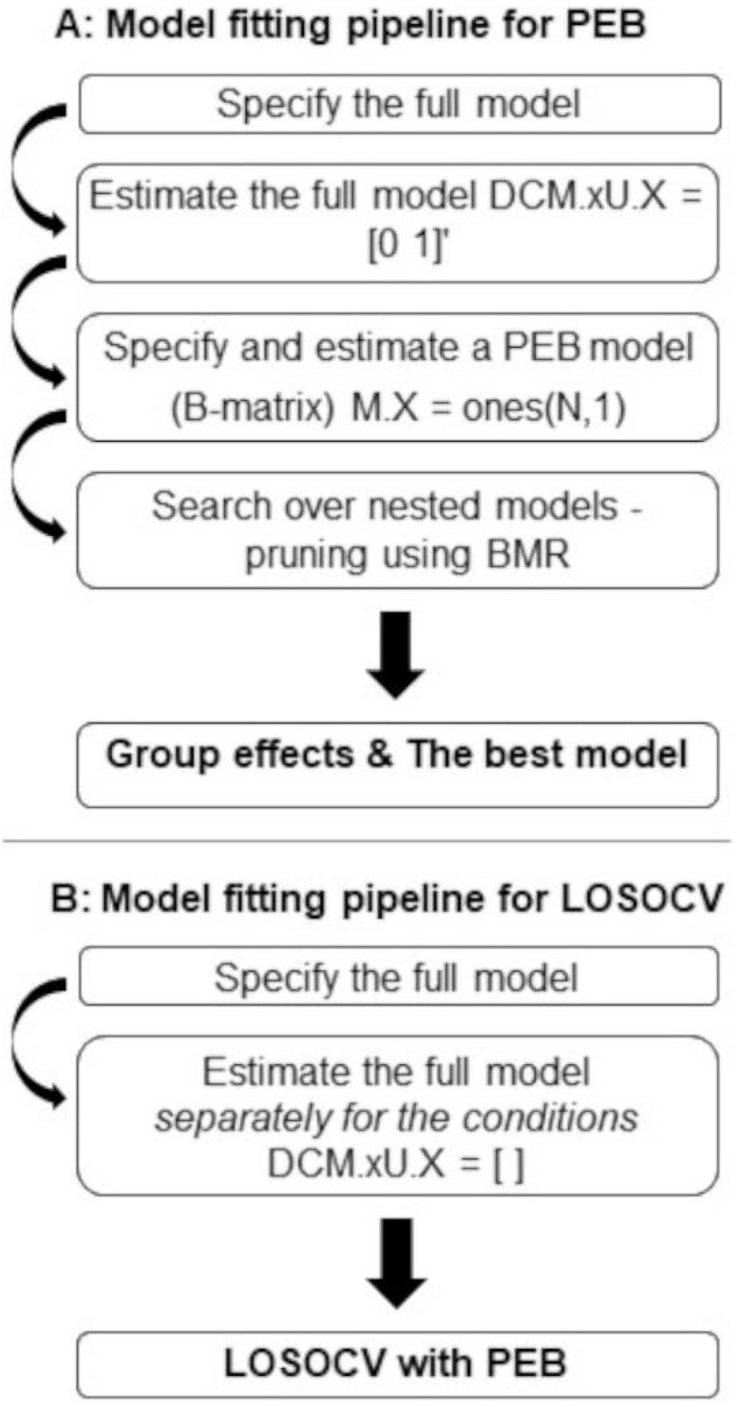
Modelling pipelines. **A.** The pipeline for inverting the DCM models in terms of changes in connectivity that explain the differences in the empirical data observed in LOC as compared to wakeful consciousness baseline. The DCM model inversion was followed by PEB modelling with BMR to find the most parsimonious model and the modulatory effects on the group-level effective connectivity. **B.** The pipeline for inverting the DCM models separately for individual states of consciousness. This was done as a prerequisite for the LOSOCV classification with PEB modelling.

### 2.5 Parametric Empirical Bayes

In DCM, a variational Bayesian scheme called Variational Laplace is used to approximate the conditional or posterior density over the parameters given by the model inversion process, by maximizing a lower bound (the negative free energy) on the log-evidence (Friston et al., 2007). The Parametric Empirical Bayes (PEB) framework is a relatively recent supplement to the DCM procedure used, for example, to infer the commonalities and differences across subjects (Friston et al., 2016). Briefly, the subject-specific parameters of interest (here, effective connectivity between nodes in a DCM model) are taken to the group-level and modelled using a General Linear Model (GLM), partitioning the between-subject variability into designed effects and unexplained random effects captured by the covariance component. The focus is on using Bayesian model reduction (BMR) – a particularly efficient form of Bayesian model selection (BMS) – to enable inversion of multiple models of a single dataset and a single hierarchical Bayesian model of multiple datasets that conveys both the estimated connection strengths and their uncertainty (posterior covariance). As such, it is argued that hypotheses about commonalities and differences across subjects can be tested with more precise parameter estimates than with traditional frequentist comparisons (Friston et al., 2016).

A particular advantage of PEB is that as part of the BMR process – when no strong a priori hypotheses about the model structure exist, as in the present study – a greedy search can be used to compare the negative free energies for the reduced models, iteratively discarding parameters that do not contribute to the free energy (originally ‘post-hoc DCM analysis’, Friston & Penny, 2011; Rosa, Friston & Penny, 2012). The procedure stops when discarding any parameters starts to decrease the negative free energy, returning the model that most effectively trades-off goodness of fit and model complexity in explaining the data. Last, a Bayesian Model Average (BMA) is calculated over the best 256 models weighted by their model evidence (from the final iteration of the greedy search). For each connection, a posterior probability for the connection being present vs. absent is calculated by comparing evidence from all the models in which the parameter is switched on versus all the models in which it is switched off. Here, we applied a threshold of >.99 posterior probability, in other words, connections with over .99 posterior probability were retained.

For the DCMs that were fitted to the contrast between two states of consciousness using the procedure described in the previous section, we used PEB for second-level comparisons and Bayesian model reduction to find the most parsimonious model that explained the contrast by pruning away redundant connections. The focus was explicitly on the group-level comparison of the connectivity modulations (B-matrix). The whole sequence of steps is summarized in figure 3A.

### 2.6 Leave-one-out cross-validation paradigm

As a crucial form of validation of our modelling framework, we investigated which network connections are predictive of the state of consciousness in unseen data. We adapted a standard approach in computational statistics, leave-one-subject-out cross-validation (LOSOCV; spm_dcm_loo.m). Here, we iteratively fitted a multivariate linear model (as described in detail in Friston et al., 2016) to provide the posterior predictive density over connectivity changes, which was then used to evaluate the posterior belief of the explanatory variable for the left-out participant: in the present case, the probability of the consciousness state-class membership.

To conduct LOSOCV analysis, the DCM models were now fitted to each state of consciousness separately, as shown in the procedure visualised in figure 3B. To cross-validate a fitted DCM model, both datasets from one participant were left-out each time *before* conducting PEB for the training data set, and the optimised empirical priors were then used to predict the state of consciousness (behavioural responsiveness/LOC) to which the datasets from the left-out participant belonged (see Friston et al., 2016 for details). This procedure, repeated for each participant, generated probabilities of state affiliation, which were used to calculate the Receiver Operating Characteristic (ROC) curves and Area Under the Curve (AUC) values with 95% point-wise confidence bounds across the cross-validation runs (see MATLAB `perfcurve`). In addition, the corresponding binary classification accuracy was calculated as the sum of true positives and true negatives divided by the sum of all assigned categories, i.e. (TP+TN) / (TP+TN+FP+FN), where TP = true positive, TN = true negative, FP = false positive, and FN = false negative.

We first estimated LOSOCV metrics for all connections in all models. Next, LOSOCV metrics of subsets of hypothesis-driven connections were tested; the connections preserved by BMR were divided into frontal, parietal, frontoparietal, and between-RSN subsets, based on the anatomical location of the connected nodes. The rationale was to investigate where in the brain the most consistent inter-subject-level effects were located, in addition to the largest effect sizes identified by the PEB analysis.

Finally, we extended our validation of the DCM models by introducing a more difficult classification problem: we used the DCM parameters from responsiveness and LOC for training, and then tested them on unseen data collected during the post-drug recovery state of each subject (recovery state prediction). Again during training, both datasets (behavioural responsiveness/LOC) from one participant were left-out each time *before* conducting PEB, and the optimised empirical priors were then used to predict the state of consciousness to which the recovery-dataset from the left-out participant belonged. We hypothesised that if our modelled effects are valid, it should classify the recovery state as behavioural responsiveness rather than LOC - even though recovery is not identical to normal wakeful responsiveness, it is clearly closer to normal responsiveness than LOC. Here, we used recall - as calculated by (true positive) / (true positive + false positive) - and mean posterior probability for responsiveness to quantify classification performance. The 95% CIs were calculated over the posterior probabilities using a simple approximation for the unbiased sample standard deviation (Gurland & Tripathi, 1971).

## 3. Results

### 3.1 Dynamic causal modeling and parametric empirical Bayes

Our goal was to investigate the effective connectivity modulations caused by anaesthesia-induced loss of consciousness on three resting state networks together and separately. We modelled time-series recorded from two states of consciousness – wakeful behavioural responsiveness and loss of consciousness (LOC) – with DCM for CSD at a single-subject level, followed by PEB at the group-level. In doing so, we estimated the change in effective connectivity with RSNs during LOC, relative to behavioural responsiveness before anaesthesia. For the DMN, we estimated 12 inter-node connections, and for both SAL and CEN 16 connections. With LAR, in addition to including all the connections in each RSN, additional connections were specified to model the modulatory effects of anaesthesia on between-RSN connections, increasing the estimated inter-node connections to fifty.

Following the inversion of the second-level PEB model, a greedy search was implemented to prune away connections that did not contribute significantly to the free energy using BMR. This procedure was performed for LAR and for all the three resting state networks separately. The most parsimonious model (A) and estimated log scaling parameters (B) for LAR, DMN, SAL, and CEN are shown in figures 4–7, respectively. Here, we applied a threshold of >.99 for the posterior probability; in other words, connections that were pruned by BMR and connections with lower than .99 posterior probability with their respective log scaling parameter are faded out (figures 4B–7B).

**Figure 4.**
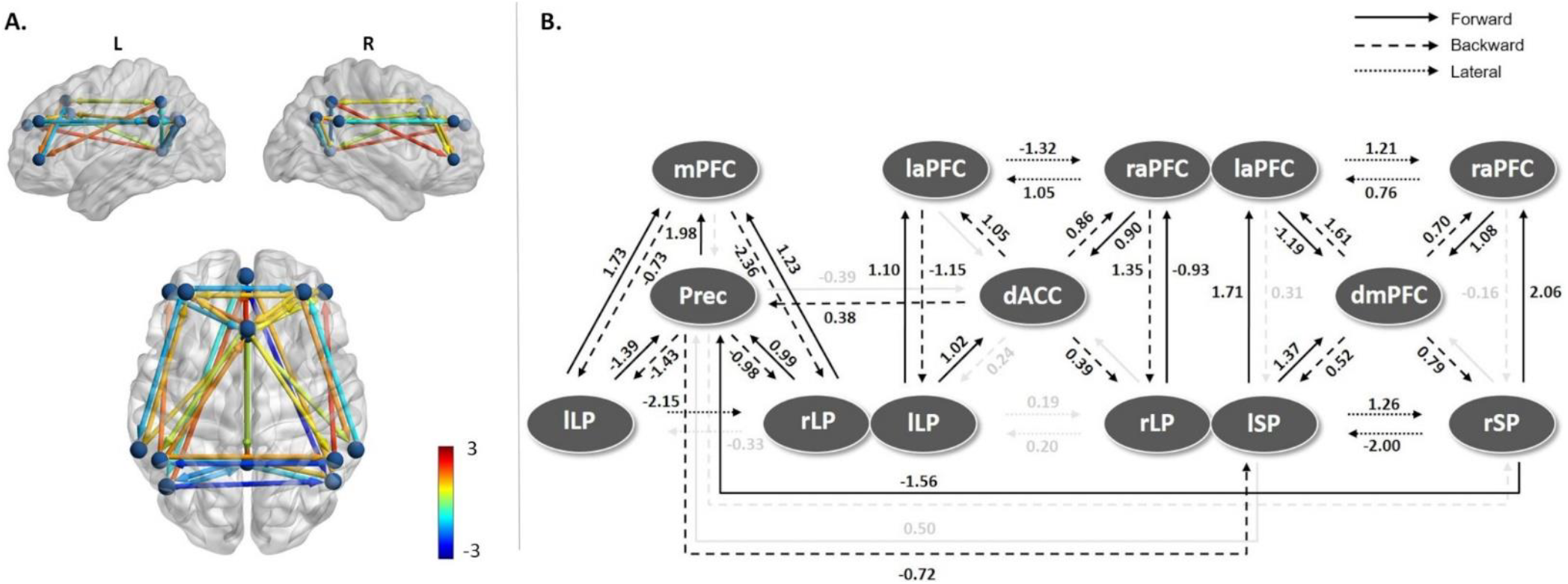
Estimated model parameters for LAR. **A.** Effective connectivity modulations on the most parsimonious LAR model. 5 connections were pruned away by BMR and a further 8 had lower than .99 posterior probability of being present. Colour shows modulation strength and direction. **B.** The log scaling parameters for the connections in the large model after BMR and BMA. Connections that were pruned by BMR and connections with lower than .99 posterior probability with their respective log scaling parameter are faded out.

Of the fifty connections in the large model (figure 4), five were pruned away by BMR. The results indicate that typically effective connectivity decreased going from behavioural responsiveness to LOC between nodes in the DMN, with parietal connections showing consistent and large decreases. Similarly, between-RSN parietal connections linking DMN and CEN also decreased. Backward connections between the dACC and PCC/precuneus, linking the DMN and SAL, increased slightly. A clear majority of connections forming the SAL and CEN networks increased.

On inverting the DMN separately (figure 5), we found that no connections were pruned away by BMR. In other words, all of the effective connectivity in the DMN was modulated by the loss of consciousness. In particular, forward connectivity to and from PCC/precuneus largely decreased, whereas direct parietofrontal forward connectivity from lateral parietal cortices to the medial prefrontal cortex was increased. Backward connectivity between all the sources was increased.

**Figure 5.**
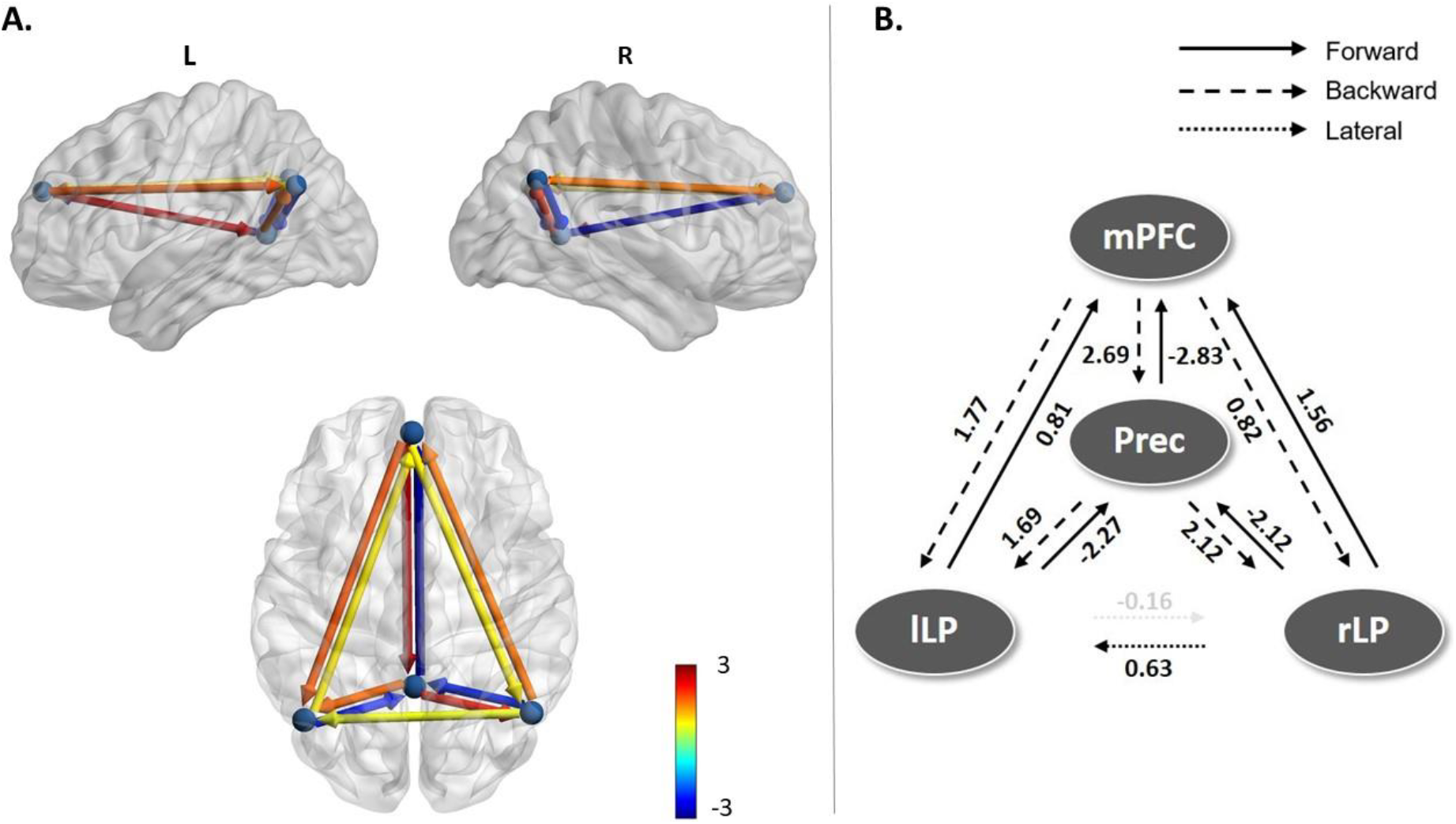
Estimated model parameters for DMN. **A.** Effective connectivity modulations on the most parsimonious DMN model. Colour of connections show strength and direction of modulation. None of the connections were pruned away, and only one connection had lower than .99 posterior probability. **B.** The log scaling parameters for the connections in DMN after BMR and BMA. The below-threshold posterior probability connection with its corresponding log scaling parameter is faded out.

In contrast, seven connections out of 16 were pruned away from the full SAL model when it was inverted separately (figure 6). These consisted of all but one lateral connections between both, the lateral prefrontal nodes and lateral parietal nodes, and all but one backward connection originating from the dACC. The strength of change in connectivity within the SAL was lower than in DMN, and all but one of the retained connections showed an increase in strength when losing consciousness.

**Figure 6.**
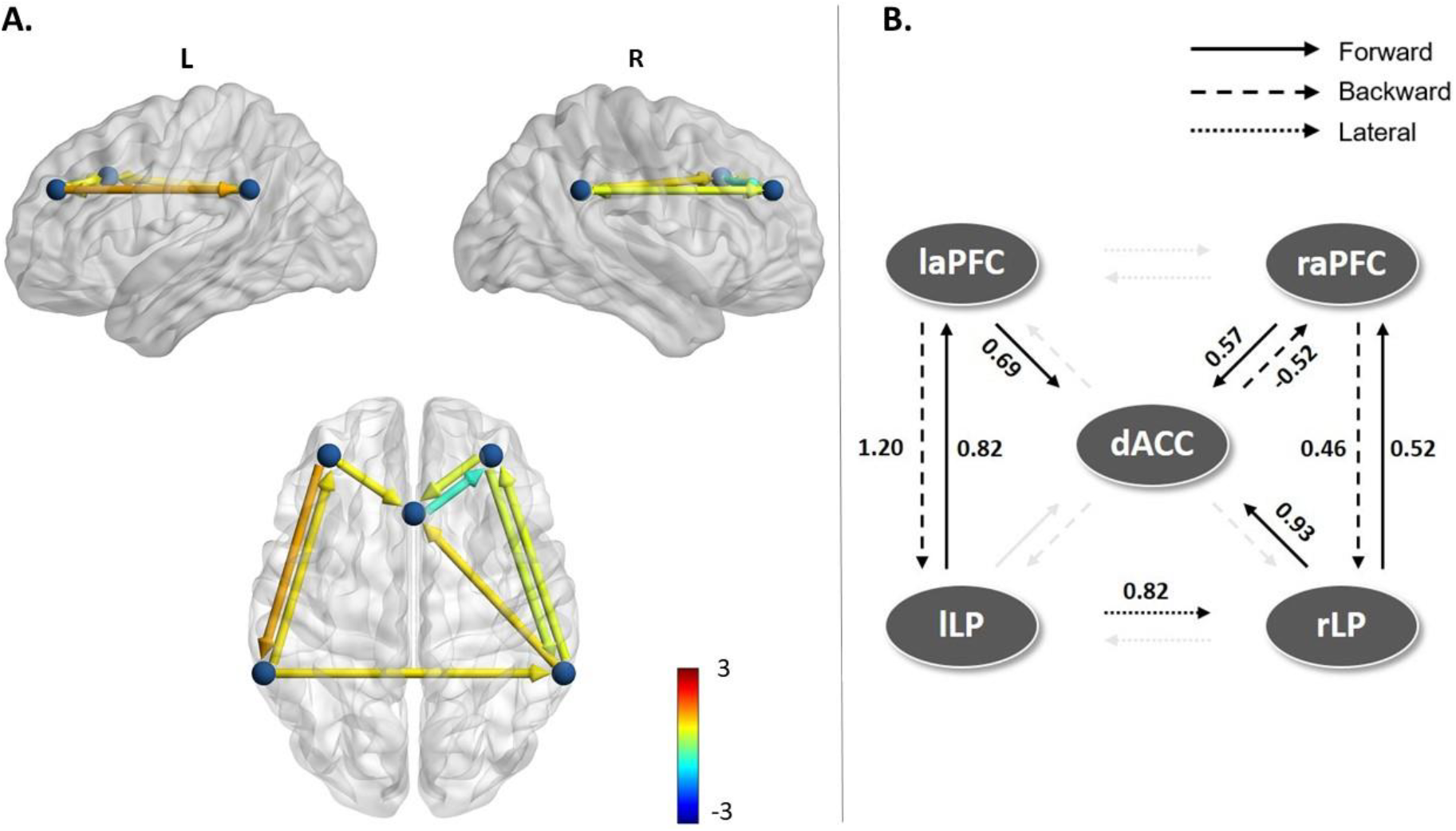
Estimated model parameters for SAL. **A.** Effective connectivity modulations on the most parsimonious model for SAL. 7 connections were pruned by BMR. **B.** The log scaling parameters for the connections in SAL. Several connections were pruned away (faded out). The retained connections were almost all positive modulations, but smaller in strength than in the DMN.

When inverting the CEN separately, two connections were pruned away (figure 7). Most of the retained connections showed a small increase in strength, with the largest effects in frontoparietal connections from the dmPFC to the left superior parietal cortex. Further, right hemisphere frontoparietal connections showed more modulatory changes than left hemisphere connections.

**Figure 7.**
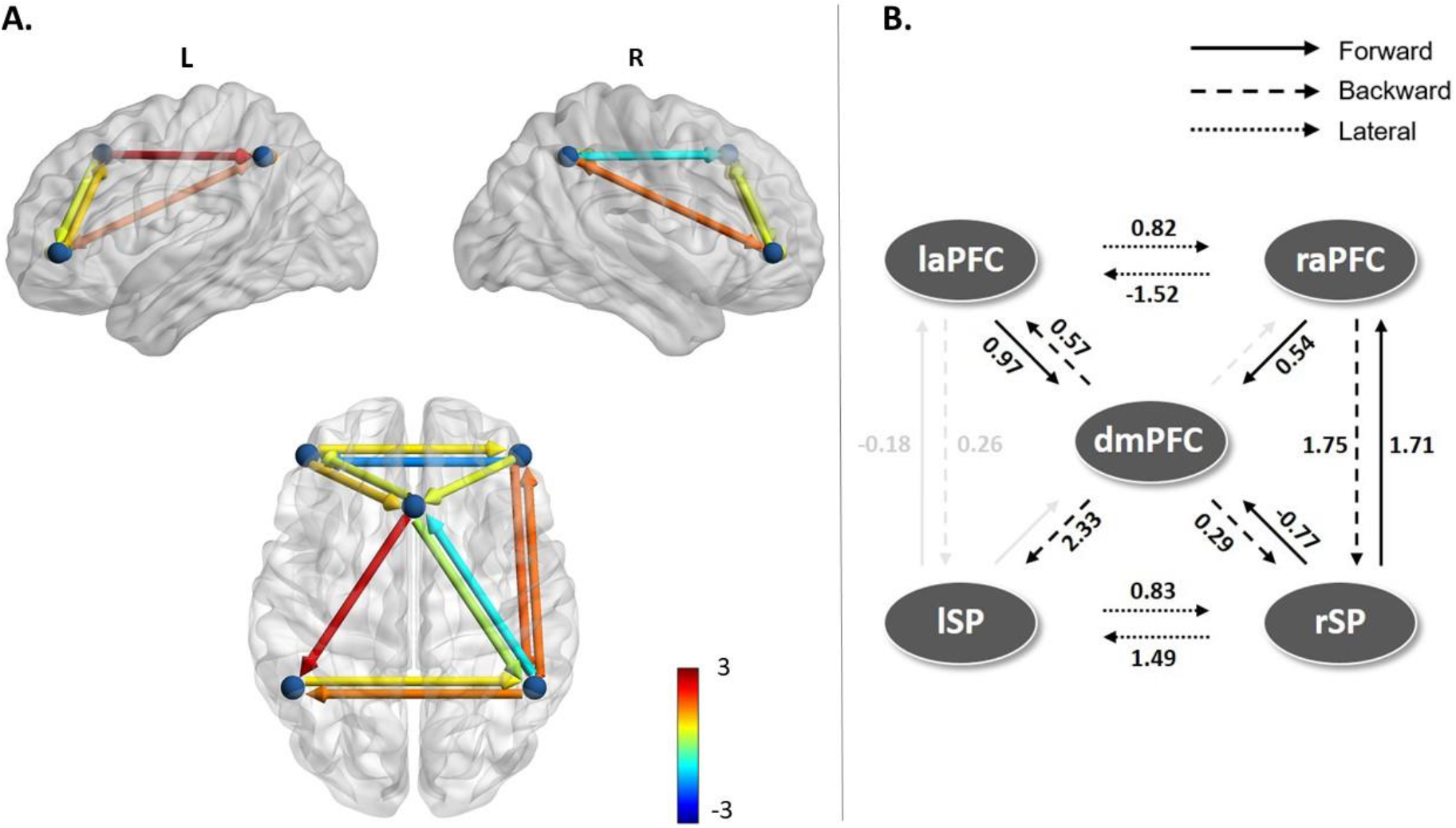
Estimated model parameters for CEN. **A.** Effective connectivity modulations on the most parsimonious model for CEN. 2 connections were redundant in addition to 2 connections having lower than .99 posterior probability for being switched on. **B.** The log scaling parameters for the connections in CEN. Pruned connections and low posterior probability connections with the corresponding log scaling parameters are faded out. Effects on the remaining connections were almost all positive modulations, with strengths in-between those observed in the SAL and DMN.

### 3.2 Leave-one-subject-out cross-validation

To conduct LOSOCV, the DCM models were inverted again, this time for each state of consciousness in each subject separately. With the states modelled separately, PEB was conducted repeatedly (on the training set in each cross-validation run) alongside LOSOCV analysis to generate AUC values (see Methods). The AUC/ROC values for all full models are shown in figure 8A, and table 2 shows all tested AUC values with accuracy for all tested sets of connections. The results indicate that leave-one-subject-out cross-validated predictions based on the LAR and SAL models had accuracy significantly different from chance, i.e. with the lower bound of the 95% CI of the AUC above chance. However, for predictions based on the DMN and CEN, the lower bound of the 95% CI of the predictions did not exceed chance.

**Figure 8.**
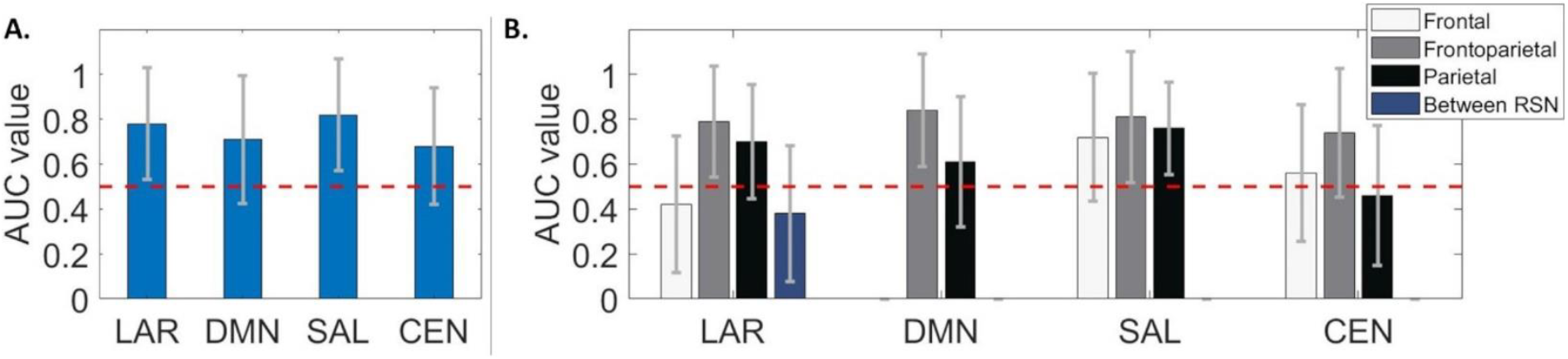
The AUC values for classifying the state of consciousness in LOSOCV paradigm. **A.** For the full models, only predictions based on LAR and SAL performed statistically better than chance (red dashed line), with classifications based on the connections in SAL reaching the overall best prediction. The error bars represent the 95% point-wise CI calculated using leave-one-out cross-validation for both A and B (MATLAB `perfcurve`). **B.** AUC values for hypothesis-driven connections for all models in LOSOCV paradigm. The DMN is missing frontal connections as it had only one anterior node. Best prediction performance was obtained with frontoparietal connections in LAR, DMN, and SAL. Further, predictions based on posterior SAL connections reached statistical significance.

**Table 2.**
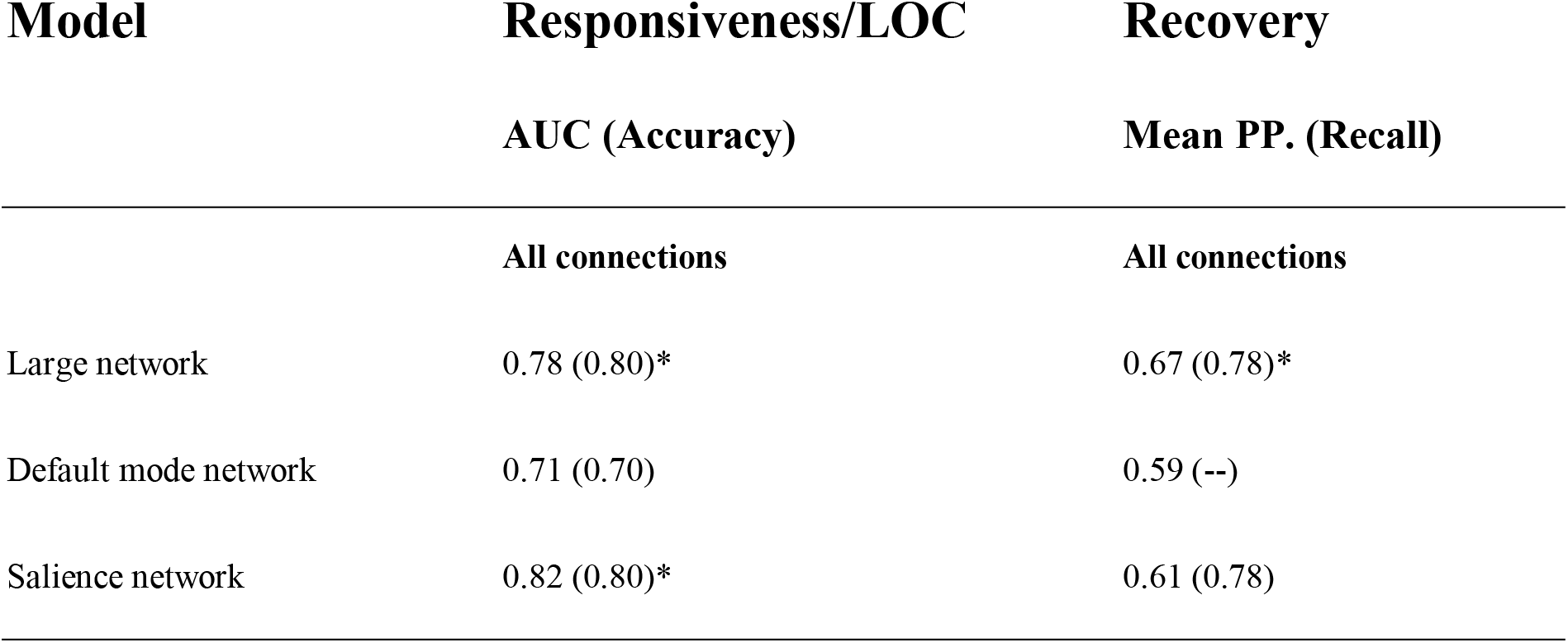
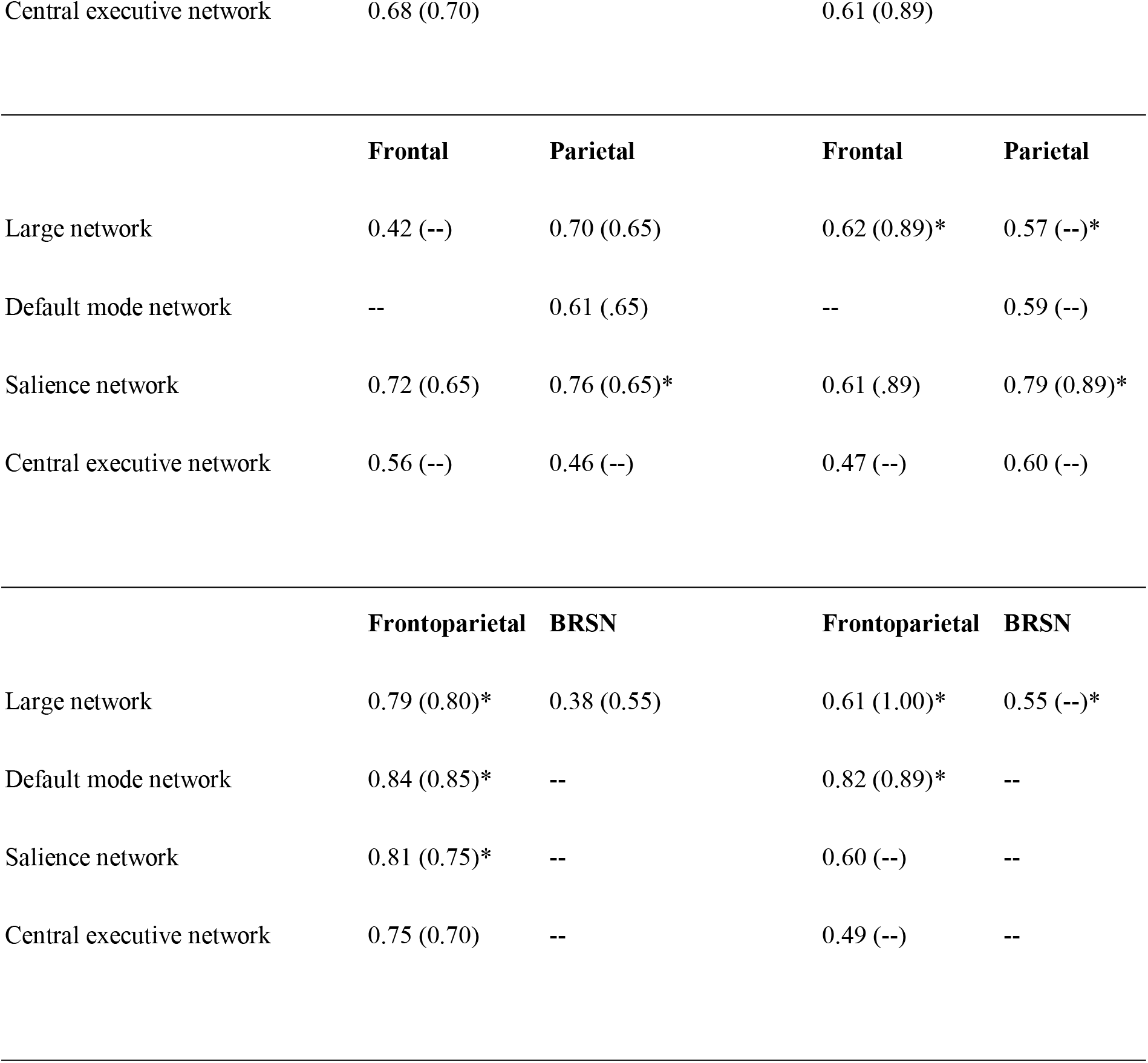
AUC (accuracy) values calculated with LOSOCV, and mean posterior probabilities (recall) in the recovery state, for all connections, all hypothesis-driven connection subsets (frontal, parietal, frontoparietal, and between-RSN connections), and all models. No values are given if no such connection-subsets exist for the model. Accuracy/recall values were not calculated for connection subsets with performance close to chance (between 0.4 - 0.6). * indicates significance estimated at 95% confidence intervals in both AUC and posterior probability.

| Model | Responsiveness/LOC | Recovery |
| --- | --- | --- |
|  | AUC (Accuracy) | Mean PP. (Recall) |
|  | All connections | All connections |
| Large network | 0.78 (0.80)* | 0.67 (0.78)* |
| Default mode network | 0.71 (0.70) | 0.59 (--) |
| Salience network | 0.82 (0.80)* | 0.61 (0.78) |

|  |  |  |  |  |  |
| --- | --- | --- | --- | --- | --- |
| 676 | Central executive network | 0.68 (0.70) |  | 0.61 (0.89) |  |
| 677 |  |  |  |  |  |
| 678 |  | **Frontal** | **Parietal** | **Frontal** | **Parietal** |
| 679 | Large network | 0.42 (--) | 0.70 (0.65) | 0.62 (0.89)* | 0.57 (--)* |
| 680 | Default mode network | -- | 0.61 (.65) | -- | 0.59 (--) |
| 681 | Salience network | 0.72 (0.65) | 0.76 (0.65)* | 0.61 (.89) | 0.79 (0.89)* |
| 682 | Central executive network | 0.56 (--) | 0.46 (--) | 0.47 (--) | 0.60 (--) |
| 683 |  |  |  |  |  |
| 684 |  | **Frontoparietal** | **BRSN** | **Frontoparietal** | **BRSN** |
| 685 | Large network | 0.79 (0.80)* | 0.38 (0.55) | 0.61 (1.00)* | 0.55 (--)* |
| 686 | Default mode network | 0.84 (0.85)* | -- | 0.82 (0.89)* | -- |
| 687 | Salience network | 0.81 (0.75)* | -- | 0.60 (--) | -- |
| 688 | Central executive network | 0.75 (0.70) | -- | 0.49 (--) | -- |
| 689 |  |  |  |  |  |
| 690 |  |  |  |  |  |

To understand whether specific connections within cortical brain networks were driving changes in consciousness, we evaluated the predictive power of four different hypothesis-driven subsets of connections – frontal, parietal, frontoparietal, or between-RSN – to predict the two states of consciousness in left-out subjects. As shown in figure 8B, frontoparietal connectivity in LAR, DMN, and SAL produced the best predictions of the state of consciousness with LOSOCV. Further, the posterior subset in the SAL performed statistically better than chance. None of the subsets in the CEN reached statistical significance.

Finally, the predictive power of these RSN connectivity subsets were tested in a more difficult classification problem: each model subset was trained on behavioural responsiveness and LOC, and then tested on the previously unseen ‘recovery’ state, the data which was collected after the participant regained consciousness. In figure 9A and B each data point represents one participant. Figure 9A shows the mean posterior probabilities of the recovery state being correctly classified as behavioural responsiveness when using all connections in a model as predictors. Figure 9B shows the same results for the frontal, parietal, frontoparietal, and between-RSN connections as predictors. When predicting with all connections, only classifications based on all connections in LAR performed significantly better than chance. With the hypothesis-driven subsets of connections, frontoparietal connectivity within the DMN generalised best to the recovery state. Only one other subset – parietal connections in SAL – performed significantly better than chance, and almost as well as frontoparietal DMN connectivity (.82 vs. .79 posterior probability). All subsets with LAR performed statistically better than chance, however, with poor mean posterior probability values in comparison to DMN frontoparietal and SAL parietal connections. Table 2 shows the mean posterior probabilities and the corresponding recall values for all the tested connection sets and for all models. We verified that the predictive accuracy (of the unseen recovery state) was not driven by subject effects or bias, as evident in the individual posterior probabilities plotted in figures 9C and 9D.

**Figure 9.**
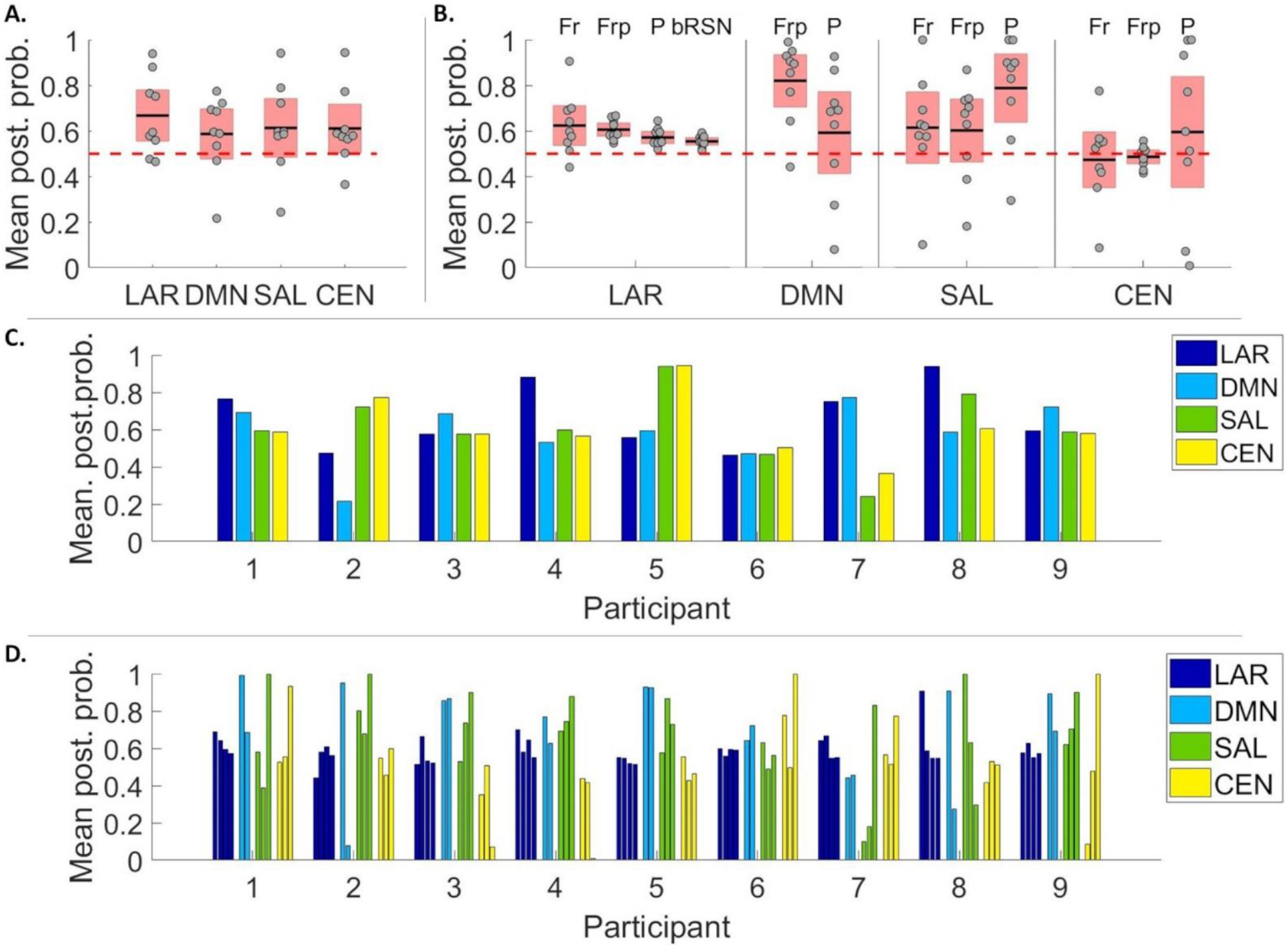
Mean posterior probabilities for prediction of recovery data. On panels A and B the individual data points represent individual participants. **A.** Predictions based on all connections in LAR performed better than chance (red dashed line). Data points representing participants are laid over a 1.96 SEM (95% confidence interval over posterior probabilities) in red with the black lines marking the mean. **B.** Mean posterior probabilities for hypothesis-driven connection subsets of all models in the recovery state: top labels refer to frontal (Fr), frontoparietal (Frp), parietal (P), and between-RSN (bRSN) connections. DMN frontoparietal connectivity had the best performance across all sets and all models. Parietal connections in SAL performed statistically better than chance but with lower posterior probability value in comparison to DMN frontoparitetal connections. All subsets with LAR performed statistically better than chance, however, with poor posterior probability values in comparison to DMN frontoparietal and SAL parietal connections. **C-D.** Posterior probabilities predicted for individual datasets, based on all connections (C) and on hypothesis-driven subsets (D). In Panel D, the individual bars depict different connection subsets: frontal, frontoparietal, parietal, and between-RSN in LAR, frontoparietal and parietal in DMN, and frontal, frontoparietal, and parietal in SAL and CEN.

## 4. Discussion

We computationally evaluated the evidence for the posterior hot zone theory of consciousness by modelling the relative contributions of three resting state networks (DMN, SAL, and CEN) for propofol-induced LOC. Using the recently introduced PEB framework, we characterised modulations in effective connectivity accompanying the loss of consciousness within and between these key RSNs. We found a selective breakdown of posterior parietal and medial feedforward frontoparietal connectivity within the DMN, and of parietal inter-network connectivity linking DMN and CEN. These results contribute to the current understanding of anaesthetic-induced LOC, and more generally to the discussion of whether the neural correlates of consciousness have an anterior contribution (Del Cul, Dehaene, Reyes, Bravo, & Slachevsky, 2009), are predominantly frontoparietal (Bor & Seth, 2012; Chennu et al., 2014; Chennu, O’Connor, Adapa, Menon, & Bekinschtein, 2016; Laureys & Schiff, 2012), or posterior (Koch et al., 2016; Koch et al., 2016b; Siclari et al., 2017).

We used a novel DCM-based cross-validation to establish the predictive validity of our models, addressing an issue commonly present in DCM studies, including previous consciousness-related DCM studies - that the best model identified by BMS is only the best model among the models tested. Significant generalisation performance with cross-validation increases the level of confidence we can ascribe to our results. This analysis highlighted that frontoparietal effective connectivity consistently generated accurate predictions of individual states of consciousness. Furthermore, we demonstrated generalisation of this predictive power by showing that effective frontoparietal connectivity within the DMN and parietal connectivity within the SAL predicted the state of consciousness in unseen data from the post-anaesthetic recovery state.

With the large model combining all 3 RSNs, we observed consistent and wide-spread decreases in connectivity between posterior DMN nodes and between parietal connections linking DMN and CEN (figure 4). With the individual RSNs, we observed a selective breakdown of the DMN, specifically, decreases in feedforward connectivity to and from PCC/precuneus (figure 5). It is worth highlighting that the largest decreases in effective connectivity - both when the RSNs were modelled individually and as one large network - were between nodes located within the posterior hot zone, and related specifically to PCC/precuneus – a key structure in the hot zone (Koch et al., 2016; Siclari et al., 2017). In other words, the network-level breakdown characterising the difference between behavioural responsiveness and LOC was mostly located within the parietal hot zone.

In the SAL and CEN networks, when fitted on their own, several connections were pruned away by BMR, with small increases in the majority of preserved connections; ¼ of the connections in CEN and almost half of the connections in SAL (7 out of 16) were pruned, in contrast to the DMN in which no connections were pruned (figures 6 and 7). The same pattern was present, although to a smaller degree, when the three RSNs were estimated together (LAR): fewest of the connections pruned were in the DMN, when compared with the SAL and CEN networks. This highlights the relative importance of the DMN over the SAL and CEN in explaining differences between states of consciousness and is consistent with the previous evidence from disorders of consciousness (Crone et al., 2011; Fernández-Espejo et al., 2012; Laureys, 2005; Laureys et al., 1999), anaesthesia (Boveroux et al., 2010), and sleep (Horovitz et al., 2009).

We found that PCC/precuneus-related feedforward connectivity in the DMN is impaired during LOC. This is in contrast to two previous DCM studies of propofol anaesthesia, which have suggested either selective impairments in frontoparietal feedback connectivity from dACC to PCC (Boly et al., 2012), or subcortico-cortical modulations from globus pallidus to PCC (Crone et al., 2017). However, there are major methodological differences between the present study and the previous two that could explain these different results. Firstly, the examined model space was different. Secondly, both previous studies used models with only two cortical nodes summarising activity of frontal and parietal regions. They did not implement a wide search over a large model space using BMR and instead focused on evaluating a small number of hypothesis-specific models. We adopted a broader approach to model formulation and evaluation. In doing so, we expand upon these previous results by suggesting a selective breakdown of PCC/precuneus-related forward connectivity within the DMN. Our results differed from Boly et al. (2012) even when the direct connections between dACC and PCC/precuneus were modelled (in LAR) – we found an increase in feedback connectivity from dACC to PCC/precuneus and a small, low probability decrease in feed-forward connectivity. Our results are, however, in line with previous studies showing increased frontoparietal connectivity with partial directed coherence (Maksimow et al., 2014) and with Granger Causality (Barrett et al., 2012; Nicolaou, Hourris, Alexandrou, & Georgiou, 2012) during anaesthesia.

It is noteworthy that impaired feedforward connectivity has been suggested to be the main modulation caused by propofol-anaesthesia in a recent DCM study with TMS-evoked potentials by Sanders et al. (2018). Their models consisted of 6 cortical sources (bilateral inferior occipital gyrus (IOG), bilateral dorsolateral PFC, and bilateral superior parietal lobule (SPL). They found predominantly impaired feedforward connectivity from right IOG to right SPL (specifically with theta/alpha-gamma coupling). Although they suggested that resting state activity was driven by feedback connectivity, while induced responses were driven by feedforward connectivity, it may be that restricting modulations to just two free parameters (connections) in the cortex simplifies the effects of propofol-induced LOC to the degree that they differ from estimations of more complex models.

Finally, the observed *increase* in effective connectivity between specific nodes (especially front-to-back) has been suggested previously to be due to the drug-specific effects of propofol rather than changes in states of consciousness (Långsjö et al., 2012; Maksimow et al., 2014). Hence, it may be that the relatively uniform increases in connectivity in the SAL and CEN, and the increased feedback connectivity in the DMN, were specific to propofol.

While the results of the LOSOCV cross-validation should be interpreted with caution given the limited number of participants in our study, the results indicated that, when using all connections, the above-chance prediction performance of conscious state was only obtained with LAR and SAL, with the latter performing the best (figure 8A). With smaller, hypothesis-driven subsets, we found that the frontoparietal connections provided consistently the most accurate predictions in all models except the CEN (figure 8B). When predicting the unseen state of recovery (figure 9B), frontoparietal DMN connections performed the best, followed by parietal connections in SAL. It is worth highlighting that the frontoparietal DMN and parietal SAL connections predict the state correctly, even when the state actually differs from the true training state; recovery differs from normal wakeful responsiveness not only behaviourally, but also in terms of the residual propofol in the blood. However, the participants are conscious and responsive, and thus, recovery is considered as a state clearly closer to normal wakeful responsiveness than LOC.

Taken together, our prediction results highlighted an important role for frontoparietal connections. This is perhaps not surprising, as wakeful awareness is known to recruit the DMN (Raichle & Snyder, 2007); maintaining a state of conscious responsiveness requires an interaction between the posterior hot zone (the role of which is highlighted when modelling the *change* between states) and frontal areas, mediated by the frontoparietal connections. Previous literature has suggested dynamic changes in connectivity between brain networks during cognitive control (Cocchi, Zalesky, Fornito, & Mattingley, 2013; Leech, Braga, & Sharp, 2012) and anaesthetic-induced loss of consciousness (Luppi et al. 2019). The importance of frontoparietal connections in the present study when predicting states of behavioural responsiveness – a state of higher integration than LOC – is consistent with the notion that conscious, behavioural responsiveness requires a brain-wide “global workspace” supported by the frontoparietal network (Baars, 1997; Dehaene & Changeux, 2011; Dehaene, Changeux & Christen, 2011; Mashour, Roelfsema, Changeux, & Dehaene, 2020). Hence, it is perhaps no surprise that the role of frontoparietal connections became prominent when we predicted individual states of consciousness rather than the contrast between them.

Lastly, a number of previous studies have suggested a pivotal role of subcortical structures in transitions to unconsciousness (e.g. Baker et al., 2014; Liu et al., 2013; White & Alkire, 2003). Crone et al. (2017) reported a breakdown of connectivity between the globus pallidus and posterior cingulate cortex connectivity during LOC, followed by a reversal at recovery. It remains a possibility that the effective connectivity modulations found in the present study – especially in relation to the PCC/precuneus - are driven by subcortical structures that we did not model here, given the limitations of scalp EEG signals (Goldenholz et al., 2009). It might be worthwhile to further investigate the effects of LOC with fMRI DCMs, including large-scale models combining cortical and subcortical nodes with PEB with BMR to conduct a wider exploration of the model space.

In addition to the modelling being limited only to cortico-cortical connections, some of our results are arguably propofol-specific; for example, very different alterations have been observed between propofol and ketamine (Driesen et al., 2013; Sarasso et al., 2015). It may be modelling the cortical effects of other anaesthetic agents would lead to very different sets of results. Despite using propofol as the tool to modulate the state of consciousness, we decided to model the effects using DCM and the standard neuronal model (ERP; based on the Jansen-Rit model), rather than models designed to better capture the subtle properties of the EEG spectrum during anaesthesia (see for example Bojak & Liley, 2005; Hutt & Longtin, 2010). Here, the methods were chosen based on the aim to model consciousness rather than the subtleties of anaesthesia. Lastly, as we tested only a pre-specified model space, the limitations imposed by this scope might have missed important mechanisms of conscious awareness not modelled here.

Notwithstanding these points, our results highlight a selective breakdown of inter- and intra-RSN effective connectivity in the parietal cortex, reinforcing the role of the posterior hot zone for human consciousness. However, modulations of frontoparietal connections were consistent enough to predict states in unseen data, demonstrating their causal role in maintaining behavioural responsiveness.

## Acknowledgements

We gratefully acknowledge support from the University of Kent’s High Performance Computing facility.

## Footnotes

1 We acknowledge that anaesthetic-induced loss of consciousness (LOC) may actually be anaesthetic-induced loss of behavioural responsiveness (LOBR), as e.g. volitional mental imagery or dreaming may take place during the anaesthetic state. The participants were, however, asked afterwards if they had any recall of dreams etc., which they did not report. Thus, here, we follow the typical convention in anaesthesia-literature and refer to this state as LOC.

2 Here, despite using propofol-anaesthesia to modulate the state of consciousness, our aim was to specifically model consciousness, rather than anaesthesia, and to produce results comparable with previous DCM EEG work with propofol. Thus, we chose the neural mass model according to our aims rather than using neuronal models designed to capture the subtleties of anaesthesia from the EEG spectrum (see, for example, Bojak & Liley, 2005; Hutt & Longtin, 2010).

